# Short-term coastal forest responses to a hurricane-scale freshwater and saltwater flooding experiment

**DOI:** 10.1101/2025.04.13.648622

**Authors:** Allison N. Myers-Pigg, Anya Hopple, Stephanie C. Pennington, Peter Regier, Ben Bond-Lamberty, Mia J. DiCianna, Kennedy O. Doro, Nate McDowell, Julia McElhinny, Alice Stearns, TEMPEST 1.0 Event Consortium, Nicholas D. Ward, Vanessa L. Bailey, J. Patrick Megonigal

**Author notes:** Materials and Correspondence to: Allison Myers-Pigg; J. Patrick Megonigal. These authors contributed equally to this work (Allison N. Myers-Pigg, Anya Hopple).

## Abstract

Coastal upland forests are exposed to intensifying precipitation regimes and sea level rise, increasing tree mortality and transforming these coastal forests into wetland ecosystems. Despite these well-known risks, the differing degrees to which hydrological, biogeochemical, and biological components of upland forests respond to novel salinity exposure is relatively unknown. The Terrestrial Ecosystem Manipulation to Probe the Effects of Storm Treatments (TEMPEST) experiment decouples two distinct disturbances associated with hydrological extremes: (1) flooding from heavy precipitation and (2) exposure to saline conditions from storm surge. Here we describe the immediate effects of saltwater and freshwater flooding on hydrologic, biogeochemical and vegetation ecosystem components following the first experimental ecosystem-scale flooding event. The experimental flooding treatments temporarily and significantly impacted the system’s hydrology but had subtler effects on biogeochemical and vegetation system components, suggesting that this temperate deciduous forest was resistant to a single novel flooding exposure, even if the water is saline. However, such episodic events can cause large transient shifts in conditions such as soil moisture and oxygen levels that may impact how the system responds to future perturbations. While the first TEMPEST event did not create substantial shifts in biogeochemical or vegetative processes, ecosystem level analysis of responses to experimental flooding through time will allow us to assess the impacts of flooding and salinity disturbances on the coupled above and belowground mechanisms driving coastal upland forest to wetland conversion.

## Introduction

The upland boundaries of coastal ecosystems are becoming increasingly exposed to hydrological extremes, causing ecophysiological stress that over time can dramatically transform upland forests to ‘ghost forests’ and then wetlands [1]. The movement of water across and within landscapes is a fundamental driver of such ecosystem state changes and also controls a wide variety of biological, physical, and chemical processes [2,3]. Shifts in inundation and saltwater exposure dynamics drive changes in vegetation community structures [1], net ecosystem production [4], carbon sequestration potential [5,6], and microbial activity [7]. As sea-level rise accelerates [8–10] and regimes of precipitation [11–14] and storms change [15–18], the emergent landscape reflects complex interactions between physical forcing (sea-level, precipitation, and storms), ecological processes (growth, competition, regeneration), and biogeochemical cycling (nutrient cycling, redox state). The mechanisms that lead to coastal ecosystem state change are vital to understand for Earth system models to generate meaningful predictions, particularly as the frequency, intensity, and rate of such changes are accelerating due to changing climate [19].

Most studies on the conversion of coastal upland ecosystems to wetlands are observation-based, using a space for time substitution design that is ultimately limited in its ability to test hypotheses about the mechanisms that underlie ecosystem state change. This is particularly true in the earliest stages of disturbance when the impacts are likely to occur belowground and be difficult to observe. Experimental manipulations are needed to unravel the rather complex cascade of linked hydrologic, belowground biogeochemical, and vegetative dynamics that are thought to drive the transition of upland coastal forests to ghost forests [20–22]. Thus, an ecosystem-level perspective is vital to understand the intertwined hydrologic, biogeochemical, and ecological mechanisms driving coastal upland to wetland ecosystem state change [23].

Here we consider the effects of novel flooding events for both freshwater and saline water on a coastal upland forest and hypothesize that hydrologic state variables (e.g., soil moisture and groundwater levels) will respond to the event immediately, followed by biogeochemical (e.g., soil greenhouse gases, soil porewaters), and finally vegetation (e.g., tree sap flux density) variables. Further, we hypothesize that the response of the upland forest ecosystem to freshwater flooding and saline flooding will be similar for hydrologic variables, while salinity changes from estuarine water flooding will have a greater impact on biogeochemical and vegetation variables compared to freshwater flooding. To test these hypotheses, we examine the effects of experimental flooding on a suite of hydrologic, biogeochemical, and vegetation response variables (Table 1) previously hypothesized to be mechanistically coupled to tree mortality [20] and/or soil microbial activity [21]. This study is part of a long-term field experiment [21] designed to decouple two distinct disturbances associated with extreme, hurricane-scale storm events: (1) flooding from heavy precipitation and (2) exposure to saline conditions from storm surge. By design, we did not attempt to simulate the many other impacts of hurricanes such as windthrow or sediment deposition [24]. This allows us to assess the impacts of hydrologic disturbances on the mechanisms driving coastal upland forest to wetland conversion through time, where we repeatedly flood the plots with water over the course of multiple years, mimicking both pulse and press disturbance through time (Fig 1).

**Fig 1.**
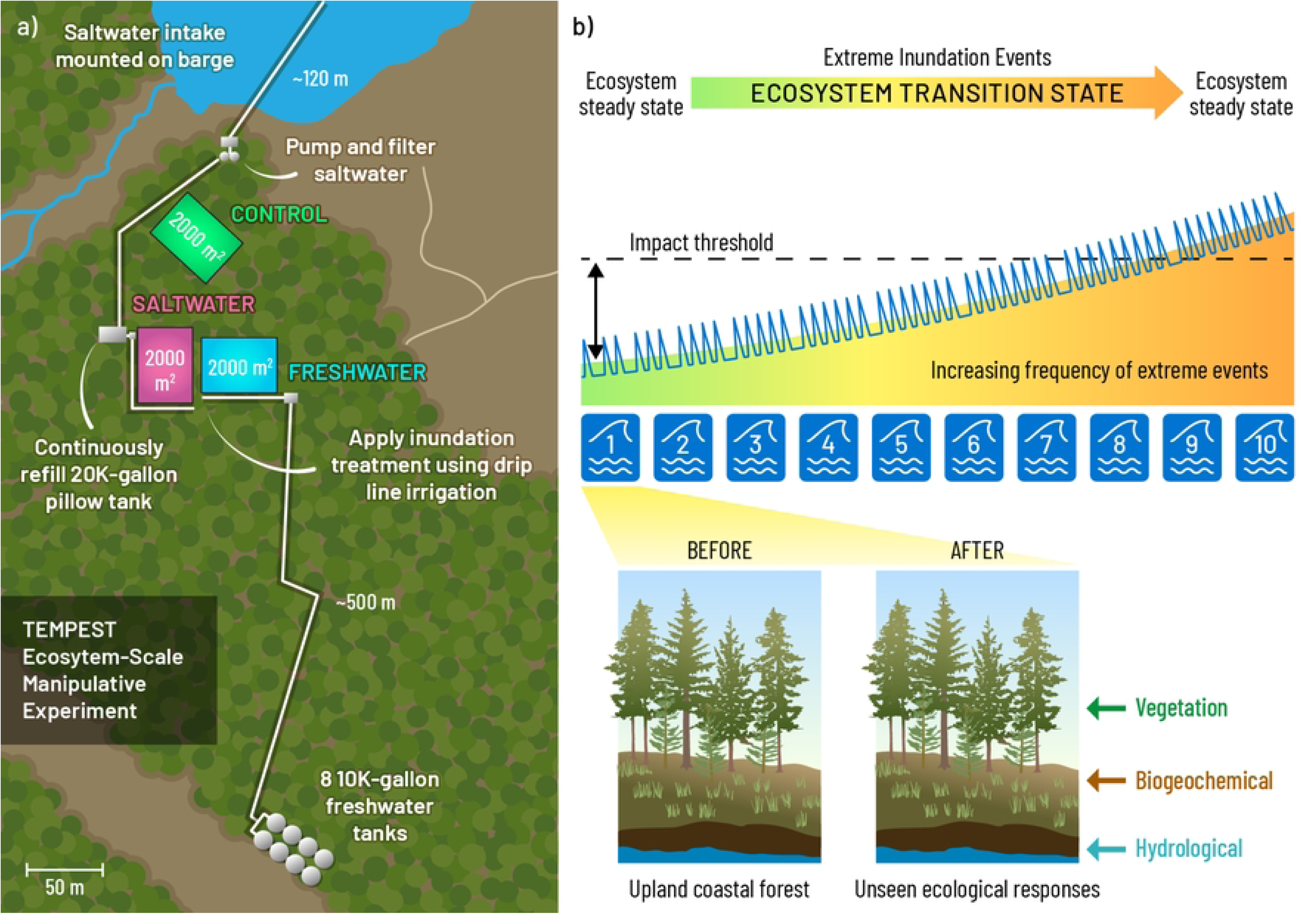
a) Site and experimental design, image modified from [21]. b) The goal of the long-term manipulation of the plots (blue line and waves) is to simulate a change in ecosystem states. Panel b is inspired by [25]. Zooming into the first event, we focus on vegetation, biogeochemical and hydrological variable changes, changes to which are not directly observable standing in the plots. Note that we employ an adaptive experimental design, where we make necessary changes to infrastructure and timing of events based on what we’ve learned in previous years.

**Table 1.** List of response variables considered in this study and their importance to coastal upland forest ecosystem functions.

| Variable | System Component | Importance |
| --- | --- | --- |
| Soil water content | Hydrological | Water availability is important for vegetation and microbes |
| Soil conductivity | Hydrological | Used as a proxy for soil salinity, which can drive mortality for species not tolerant to it |
| Groundwater depth | Hydrological | Metric of vertical connectivity between surface soils and the groundwater system |
| Groundwater conductivity | Hydrological | Metric of vertical connectivity between surface soils and the groundwater system, these values should be low in an unperturbed upland forest system |
| Soil porewater dissolved organic carbon (DOC) | Biogeochemical | Dissolved organic carbon is a function of and driver of biogeochemical processes, including microbial processing of organic matter in soils |
| Microbial methane (CH<sub>4</sub>) flux | Biogeochemical | Metric of greenhouse gas production from soils without the influence of plant |
|  |  | roots, defined here as from microbial processes. |
| Microbial carbon dioxide (CO<sub>2</sub>) flux | Biogeochemical | Metric of greenhouse gas production from soils without the influence of plant roots, defined here as from microbial processes. |
| Soil oxygen (O<sub>2</sub>) | Biogeochemical | Hypoxia or anoxic conditions can stress vegetation as well as alter microbial decomposition pathways. |
| Soil porewater spectral slope ratio | Biogeochemical | Indicator of the quality of organic matter present in soil porewaters, which impacts its ability to be degraded by microbial communities in the soil system. |
| Groundwater dissolved oxygen (O<sub>2</sub>) | Biogeochemical | Hypoxia or anoxic conditions can stress vegetation as well as alter microbial decomposition pathways. |
| Tree sap flow velocity | Vegetation | Provides insight into how the tree is functioning and its overall health, including its stress levels. |
| Root-influenced CH<sub>4</sub> flux | Vegetation | Metric of greenhouse gas production from soils, including plant roots. |
| Root CO<sub>2</sub> flux | Vegetation | Metric of greenhouse gas production from plant roots. |

## Methods

### Site description

The Terrestrial Ecosystem Manipulation to Probe the Effects of Storm Treatments (TEMPEST) experiment is set in a temperate, deciduous forest on the western shore of Chesapeake Bay in Maryland, USA (38.876°N, 76.553°W) at the Smithsonian Environmental Research Center (SERC: https://serc.si.edu/). The ∼60-year-old upland coastal forest is dominated by *Liriodendron tulipifera*, *Fagus grandifolia*, *Acer rubrum*, and *Quercus* spp., while the understory is composed of deciduous shrubs such as *Rubus phoenicolasius*, *Lindera benzoin*, and *Berberis thunbergii* and herbaceous perennials such as *Mitchella repens*, *Polygonum virginianum*, *Rhus radicans*, and *Symphyotrichum lateriflorus*. The water table is on average ∼2 m below ground and is drained by a second-order stream that flows into a brackish tidal marsh with a mean annual salinity of 10 psu and tidal range of 44 cm. The soils are well-drained fine sandy loams or sandy loams classified as Typic Hapludults [26]. More details on the soils and vegetation characteristics can be found in [21].

TEMPEST simulates extreme, ecosystem-scale freshwater and saltwater disturbance events using a novel, large-unit (2000 m^2^), un-replicated experimental design, with three 50 m × 40 m plots serving as control, freshwater, and saltwater treatments. These plots had no known prior exposure to saline conditions. A high-resolution spatiotemporal approach achieved with a grid-system strategy is used to monitor the impacts of experimental treatments on subsurface hydrology, biogeochemistry, and vegetation across these large plots in response to the simulated flooding events. Plots are designed to spatially coordinate measurements spanning the soil-plant-atmosphere continuum. Temporal patterns are captured by continuous sensor networks complemented by discrete measurements at regular bi-weekly or monthly intervals, with higher frequency immediately prior to, during, and following simulation events [27].

### Flood manipulation

Detailed methodology on the flood manipulations can be found in [21]. Briefly, intake systems draw freshwater from a municipal source and brackish water from the adjacent Rhode River estuary, which is then distributed through a network of irrigation tubing equipped with pressure-compensating emitters. The water delivery rate is just above the drainage capacity of the soil and was selected to maximize the time that the soil remains saturated while minimizing water loss by surface runoff. For each flooding event, we aim to deliver 300 m^3^ of water, ∼15 cm, to each treatment plot at an average rate of 640 L per minute (LPM) over a 10-h period, per event. For simplicity, we refer to the plot where brackish water treatment was applied as the “saltwater” plot.

### Hydrological, biogeochemical and vegetation data collection

#### Soil and groundwater in-situ sensors

Soil temperature, moisture content, and electrical conductivity (EC) were measured using TEROS 12 soil sensors (Meter Group) deployed at 5, 15, and 30 cm depths in five grid cells in each plot and deployed at 15 cm in an additional 31 grid cells in each plot (Fig S1). Soil oxygen (O_2_) was measured using Firesting optical oxygen sensors (Pyroscience, Germany) at 30 cm depth in a single grid cell in each plot installed prior to the first flood event. Groundwater depth, salinity, and dissolved O_2_ were measured by Aqua TROLL 600 multiparameter sondes (In-Situ) installed in ∼4 m deep groundwater wells in a single central grid cell in each plot. A generalized map of sensor installation locations across the experimental plots can be found in the Supporting Information (Fig S1).

All sensor data were first visually inspected for outliers or sensor malfunctions. TEROS sensor datasets (soil temperature, moisture, and EC) were linearly gap-filled for any missing data (<2% missing for every sensor, with a maximum gap of 2 hours), then binned by timestamp across all sensors for each plot. For Firesting datasets, we replaced all values < 0 with 0, which we suggest are due to small differences (minimum measured value: -0.254 mg/L) between actual 0 and the calibrated value for 0. Groundwater depths below the soil surface were calculated from pressure and well dimensions after correcting for atmospheric pressure and water density. Each of the resulting datasets contained 15-minute time-step resolution data at the individual plot scale.

#### Soil porewater

Grab samples of soil porewater chemistry were collected from 10 permanently installed lysimeters at 15 cm depth, distributed across each of the plots (Fig S1). Samples for measuring optical properties of soil porewater were pooled within a given plot due to volume limitations. Samples were field filtered immediately after collection using a 0.45 µm syringe filter (MilliporeSigma™ Millex™ Nonsterile 33 mm Syringe Filters) and stored at 4 °C until analysis. Dissolved organic carbon (DOC) was measured on filtered samples within one week of collection on a Total Organic Carbon Analyzer (Shimadzu TOC-L). DOC was measured as non-purgeable organic carbon (NPOC) via catalytic combustion after in-line acidification with 1:12 hydrochloric acid. Check standards were run every 10 samples. Data underwent additional quality control, including visual inspection of calibration curves, check standards, and sample peak shapes. Peaks were disregarded if the coefficient of variation between replicate injections was greater than 2.0%, and values were flagged when they were outside of the calibration curve and instrument detection limit ranges. Further, values were removed from subsequent analysis if dilution factors were greater than 30, blank values were greater than or equal to 25% of raw instrument sample values, or when replicate samples were greater than 25% apart. UV absorbance scans and excitation-emission matrices (EEMs) were collected simultaneously on filtered samples using an Aqualog (Horiba Scientific), with absorbance measured from 230 to 800 nm in 3 nm intervals. EEMs were collected within the same wavelength constraints and were further processed with the drEEM toolbox v. 6.0 for Matlab. Absorbance data were blank-corrected prior to exporting data in the Aqualog software. Processing of the EEMs in the drEEM toolbox included blank correction, inner filter correction, and normalization to Raman Scatter units based on daily water Raman scans collected at an excitation of 350 nm.

#### Greenhouse gases

Soil CH_4_ and CO_2_ flux measurements were taken at permanently installed collars distributed throughout the plots using an infrared gas analyzer (IRGA; LI-7810, LI-COR Inc., Lincoln, NE) attached to a 20 cm-diameter soil flux chamber accessory (LI-8201 Smart Chamber). The IRGA measured concentrations every second over a 1-min period and calculated flux based on a nonlinear regression of gas concentration in the closed chamber system over time per unit area. Two successive measurements were taken at each collar and averaged using the LI-COR SoilFluxPro software (v4).

We installed a root-exclusion experiment within each of the three experimental plots to understand the role of roots on soil CH_4_ and CO_2_ emissions [28]. Briefly, we established eight small (0.5 m^2^) subplots in each of the large experimental plots to impose two treatments replicated four times each. We trenched four of the subplots to a depth of 60 cm and lined the resulting monoliths with 45 µm mesh to exclude roots alone (i.e. root_free plots) but not mycorrhizae. The other four subplots were undisturbed control plots that were not trenched. Four additional subplots, lined with 1 µm mesh as reported in [28], were not used here. In the present study we report two response variables. We define ‘root-free fluxes’ as those attributed to microbial activity. We also report fluxes attributed to roots calculated as Flux_undisturbed_ - Flux_root_free_. For soil CO_2_ flux this quantity is root respiration, and for CH_4_ it is ‘root-influenced’ soil CH_4_ as roots do not emit meaningful amounts of CH_4_. The two treatments used for the present analysis are a subset of four treatments that constitute the full experimental design as detailed in [28].

#### Sap flow velocity

Tree transpiration was measured to assess the impacts of the experimental floods on vegetation. Sap flow velocity was measured using the thermal dissipation method (Plant Sensors, Nakara, Australia; [29]), where two 3.5-cm probes with a 2-cm sensing length were installed 10 cm apart. The top probe is heated at a constant power and the bottom is a reference probe. Sensors were installed in 18 trees per plot constituting six replicates in each of three tree species, *Acer rubrum* (red maple), *Liriodendron tulipifera* (tulip polar), and *Fagus grandifolia* (American beech) and measured every 15-minutes continuously. Sap flow velocity, in cm/hr, was calculated based on [30] (Equation 1):

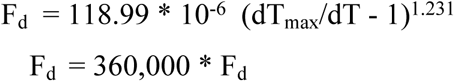

where dT is the difference in the heated and reference probes and dT_max_ is the maximum sap flow between the hours of 12:00 am and 5:00 am when sap flow is zero [29]. We only used daytime sap flow data (defined as the hours of 5:00 am and 9:00 pm) for this analysis, to reduce the influence of inter-species differences in stem capacitance on transpiration [31]. Sap flow velocity was averaged by plot.

#### Ecosystem-level analysis

Each variable was processed to include data between 20 June 2022 and 25 Jun 2022, except for porewater that used 13 Jun 2022 as a start date due to sampling frequency differences. These windows capture 48 hours before and after the experimental events. We only included data from TEROS sensors at 15 cm depth in the calculations for Equation 2 below, to draw from the largest number of sensors and spatial variation across the plots. Analytes were kept at the collection frequency for each variable. To assess changes and test our hypotheses across heterogeneous data types, data were first binned into five distinct groups: 1) Pre (pre-events, <22/06/2022 05:30 EST), 2) Mid (flood event, 22/06/2022 05:30 – 22/06/2022 14:30 EST), 3) Between Events (22/06/2022 14:30 EST – 22/06/2022 18:00 EST), 4) Rain Event (22/06/2022 18:00 – 23/06/2022 22:30 EST), 5) Post (post-events, > 23/06/2022 22:30 EST). The rain event was a natural event that occurred less than 24 hours after the end of the planned experiment. We then calculated the change in each response variable during the event as (Equation 2):

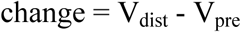

where V_pre_ is the average variable value of the Pre time period (baseline) and V_dist_ is the largest absolute value during the disturbance event – either the minimum (variable decreased during event) or maximum (variable increased during event) variable value during the Mid time period.

We calculated the minimum and maximum changes calculated per Equation 2 that was observed across all variables in the control plot. We define responses greater or less than the control plot variability as outside of natural variance and therefore a response to flooding.

#### Statistics

This is an unreplicated study at the plot level, which limits our ability to do certain statistical tests, such as a direct comparison of changes through time with simple parametric tests. As we are comparing a small window of time for this analysis (48 hours pre/post event), and due to different sampling frequencies across variables, we also could not use the before-after-control-impact (BACI) statistical framework in which the larger experiment was designed [21]. Wherever percent difference is reported, it was calculated based on plot averages across timepoints, and no statistics were performed on these percent difference values to avoid possible pseudo replication issues [32].

To test our hypotheses, we required a statistical design that assess differences in change among system components/response variables. Therefore, we adapted statistical approaches from other system-level analysis works to assess the differences among response categories (i.e. system components) using analysis of variance (ANOVA) followed by Tukey-HSD post-hoc test, when significant [33]. All statistical tests were conducted in R version 4.4.0 [34]. Datasets were tested for normality (Shapiro-Wilk’s test) and equal variance (Bartlett’s test), and we rank-normalize data to achieve normal distributions prior to statistical testing when assumptions of normality and equal variance were not met. To explore the differences in the change (immediate response during event) across the treatment plots (freshwater/saltwater) and system components (hydrological, biogeochemical, and vegetation), we conducted a two-way ANOVA (change ∼ system component x plot). To assess differences in changes observed (immediate response during event) within each system component (e.g. for the variables within the hydrological component) across all plots (control, freshwater, saltwater), we conducted a one-way ANOVA for each system component (change ∼ system component).

All data and analytical code to reproduce our results are available at https://github.com/COMPASS-DOE/tempest-system-level-analysis.

## Results

### Treatments

We delivered 263 m^3^ of freshwater and 267 m^3^ of brackish water, equivalent to ∼13 cm across the 2000 m^2^ plots over nine hours on 22 June 2022 (Fig 1; Table S1). This resulted in widespread saturation of the forest soils during the event, consistent with our system test using only freshwater on both plots the prior year [21]. Incoming water during the experiment had a conductivity of 127 ± 4 µS/cm for the freshwater plot, and 13,666 ± 1,206 µS/cm for the saltwater plot (equivalent to 7.9 ± 0.7 PSU).

Soon after the experimental flooding event, 2.8 cm of rainfall occurred over 29 hours. This natural event had a noticeable impact on all hydrologic variables, soil O_2_ (biogeochemical), and sap flow velocity (vegetation).

### Change among ecosystem components

We first examined responses to flooding for hydrological, biogeochemical, and vegetation variables as a change relative to the control plot (Fig 2). The change was different across system components (p <0.05) but showed no difference between the treatment plots (freshwater and saltwater, p = 0.557). Statistical differences across the system components (hydrological, biogeochemical, and vegetation; p < 0.01) showed a very weak interaction with plot (p = 0.094). Post-hoc tests revealed that hydrologic variable responses were different from both biogeochemical and vegetation responses (p < 0.05), with no significant difference between vegetation and biogeochemical responses in terms of change.

**Fig 2.**
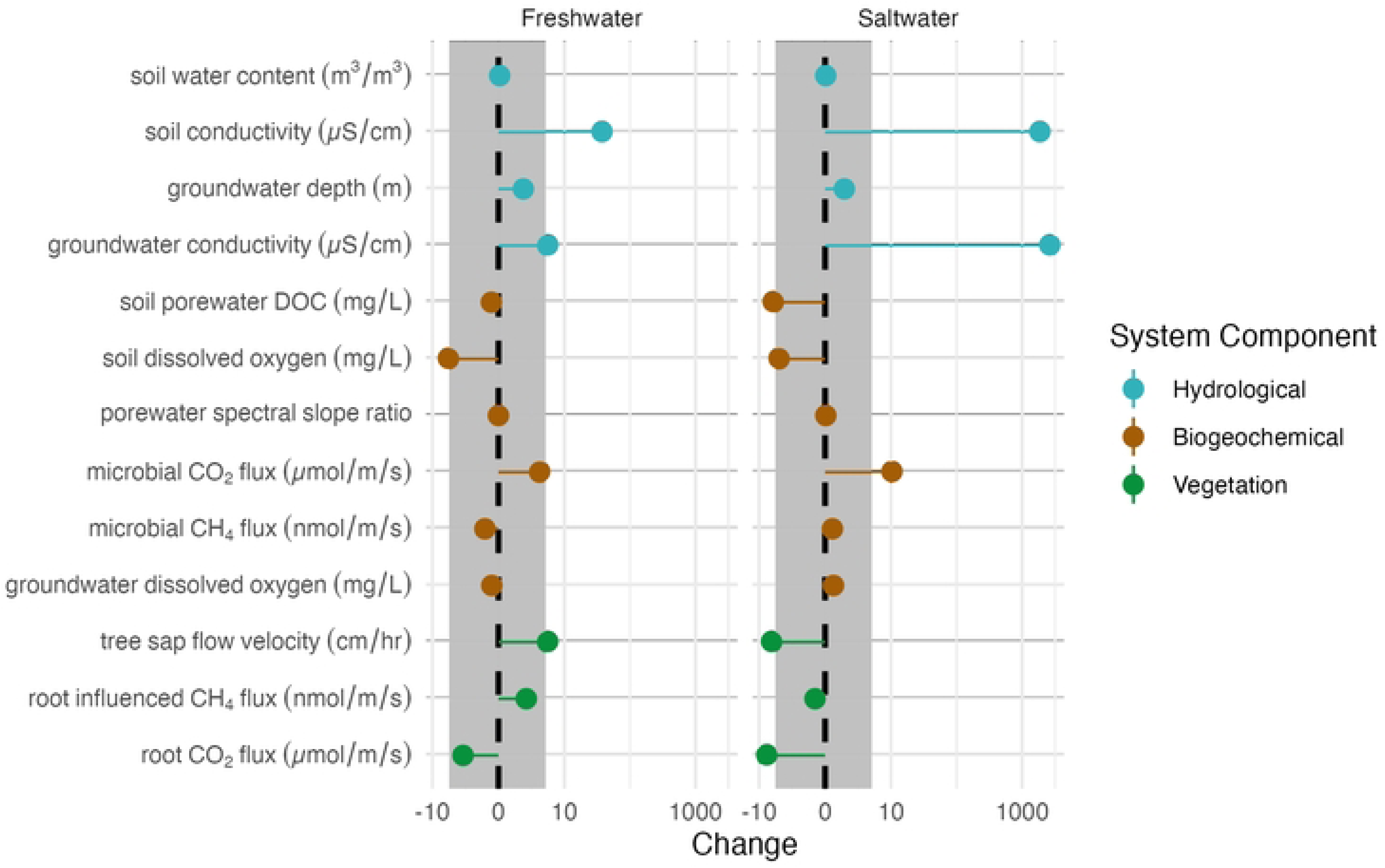
Change in response variable (note log scale on x axis) for measured biogeochemical, hydrological, and vegetation variables during the event compared to 48 hours prior. The grey shaded area indicates the change in control plot for all variables during the study period.

We found significant differences in the change from before to during the event (Equation 2) between plots for the hydrological variables (p < 0.01), which post-hoc Tukey-HSD analysis indicated was driven by differences between the saltwater and the control plot (p < 0.01) and the freshwater and the control plot (p = 0.014). No statistical differences in the change from before to during the event were observed across plots for the biogeochemical (p= 0.833) and vegetation (p=0.114) system components, although we note that subtle changes in these variables through longer time periods than we considered here may still influence ecosystem functions.

### Hydrologic responses

Hydrologic variables had the largest immediate responses to the TEMPEST event in both treatment plots. Soil water content and electrical conductivity at 15 cm depth increased in both plots during the event when compared the control plot (Figs 2 and 3). Soil water content peaked toward the end of the treatment event for both the freshwater and saltwater plots and started to decrease immediately after the end of the treatment (Fig 3). During the event, soil water content in the freshwater plot increased ∼20% from baseline, and ∼10% from baseline in the saltwater plot (Fig 3).

**Fig 3.**
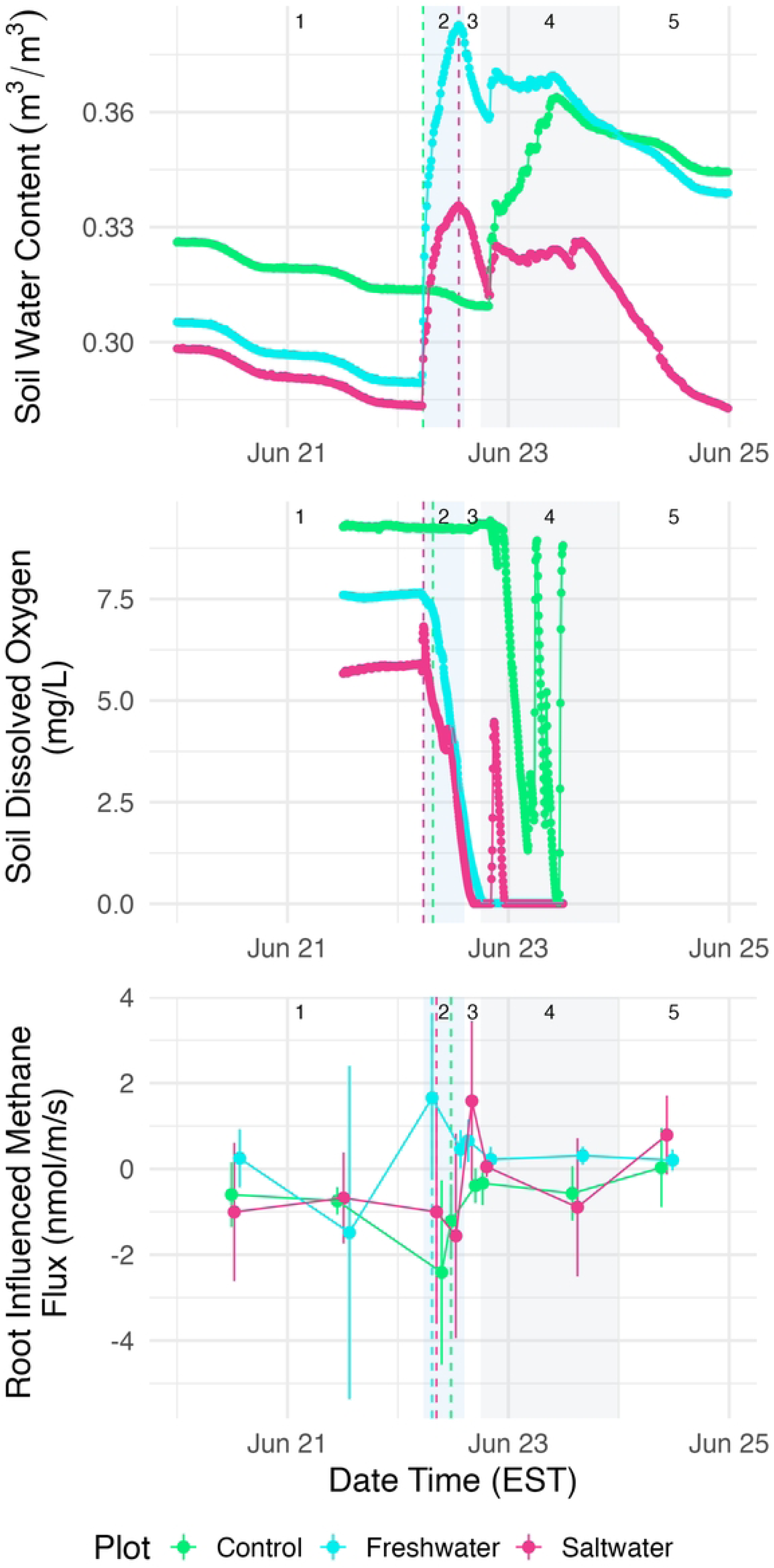
Time series of example variable responses through study period for hydrological (soil water content), biogeochemical (soil O_2_), and vegetation (root-influenced methane) system components, 48 hours before, during, and 48 hours after the event. Box 1 highlights pre flood event, box 2 is during the flooding event (blue), box 3 is after the flood event, box 4 is the rain event following the flood event (grey), and box 5 is after the rain event is over. Dashed lines in box 2 represent the Vdist used for the change calculations for each plot in the respective plot colors. Note that for soil O_2_ the freshwater and saltwater Vdist are five minutes apart, and for soil water content freshwater and saltwater Vdist are at the exact same time stamp.

An order of magnitude difference between the freshwater and saltwater plots for soil electrical conductivity at 15 cm persisted throughout the soil profile (Fig 4). Soil electrical conductivity had the greatest overall response of all variables, with an increase that exceeded the control plot variance envelope in both the freshwater and saltwater plots (Fig 1). During the flooding event, conductivity at 15 cm depth increased by ∼ 40% compared to baseline in the freshwater plot, and ∼2700% compared to baseline in the saltwater plot (Figs 2 and S2).

**Fig 4.**
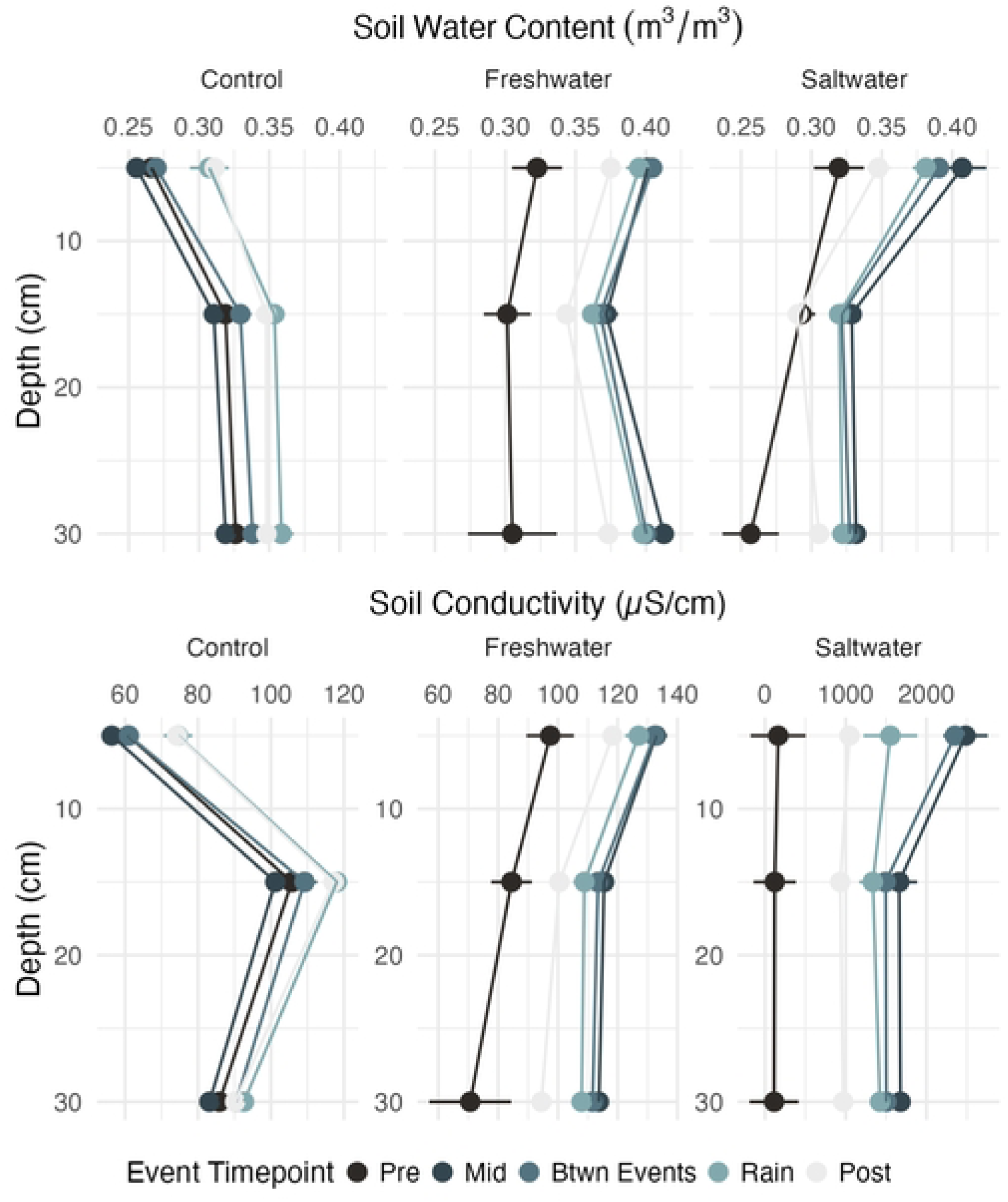
Depth profiles of a) soil water content and b) soil conductivity for the 48 hours prior to the event (darkest shade), middle of flooding event, between flooding and rain event, during the rain event, and 48 hours after the rain event (lightest shade) for control (left), freshwater (middle) and saltwater (right) plots. Note variable x axis on panel b.

Groundwater levels and salinity increased in both plots during the event (Fig S2). The groundwater level rose from a depth of ∼3 m below the soil surface to within the root zone in both the saltwater (57 cm minimum depth) and freshwater plots (29 cm), peaking in both plots shortly after water application stopped. Groundwater electrical conductivity increased outside of the control plot variance envelope for both the freshwater and saltwater plots (Fig 2).

### Biogeochemical responses

Most biogeochemical parameters monitored in the soil, porewater, and groundwater responded similarly between freshwater and saltwater treatments relative to the control plot. These responses were either large, in the same direction, and of similar magnitude (soil dissolved O_2_, microbial CO_2_ flux), or of small magnitude, with varying directions of change (soil porewater DOC, soil porewater spectral slope ratio, microbial CH_4_ flux, groundwater dissolved O_2_) (Table 2). Three biogeochemical variables had changes that were outside the control plot variance: microbial CO_2_ fluxes were higher than control variance in the saltwater plot, soil O_2_ was lower than control variance in the freshwater plot, and soil porewater DOC was lower than control variance in the saltwater plot.

**Table 2.**
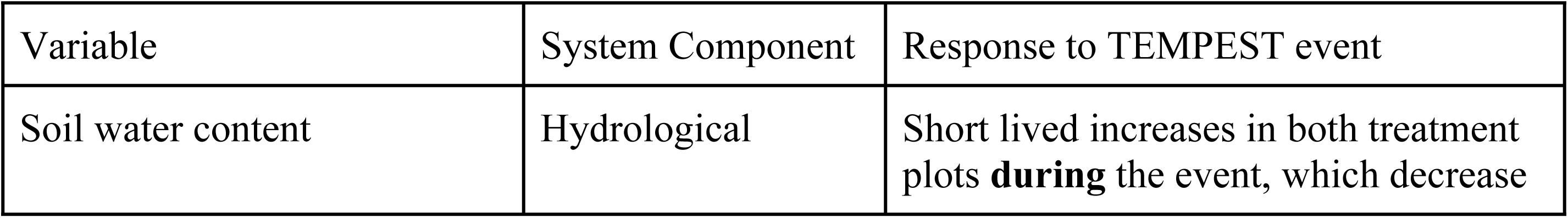
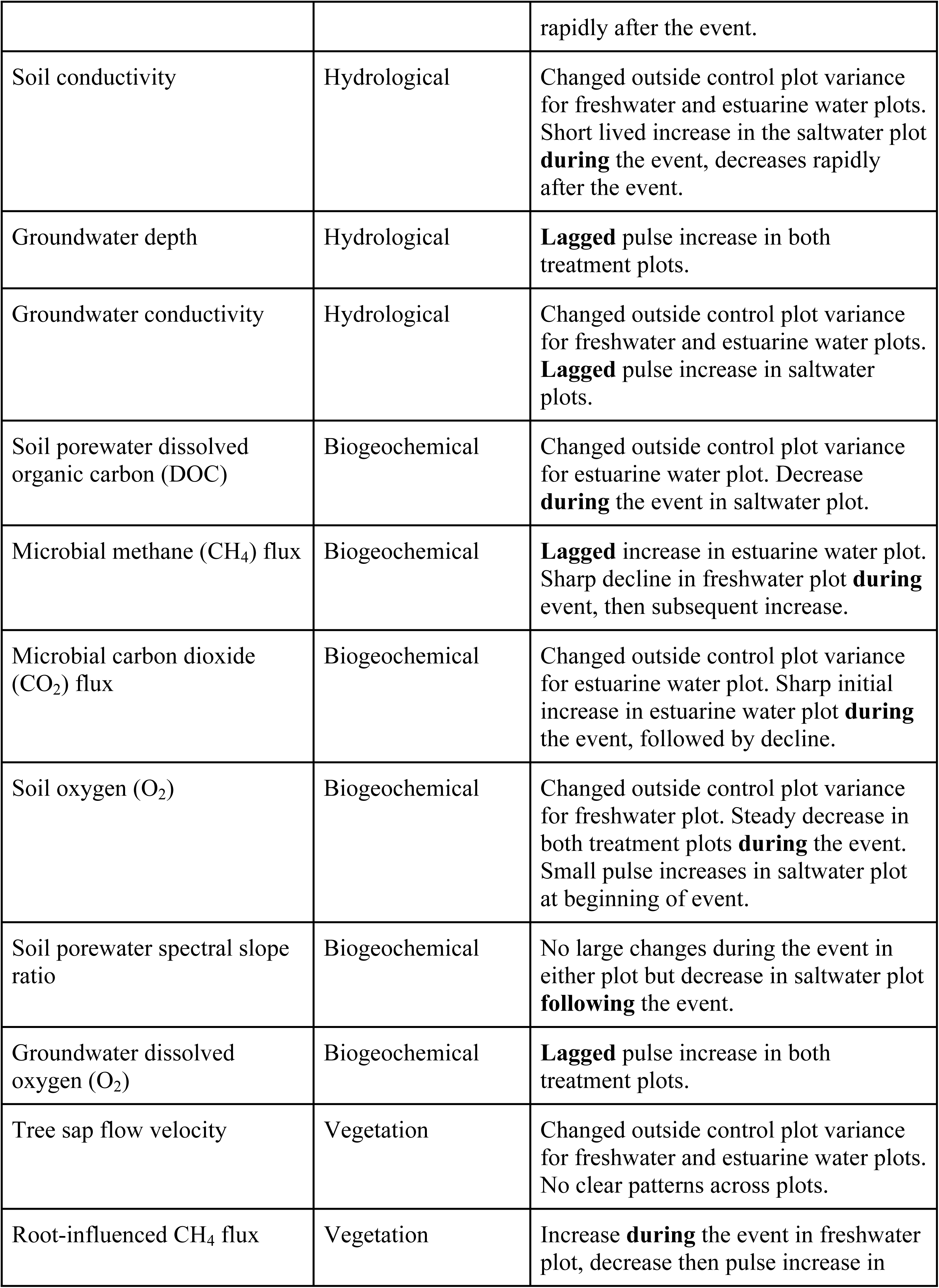
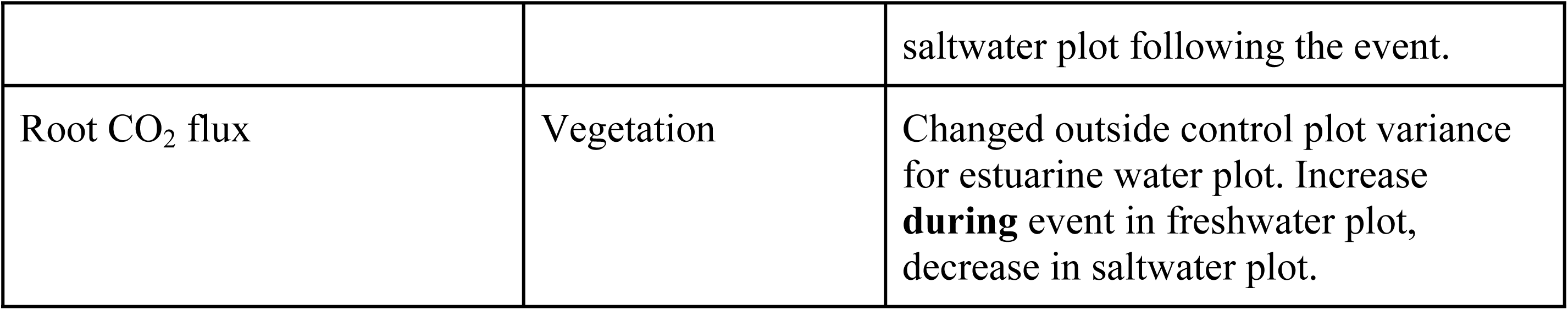
Major system responses to the first TEMPEST event for each system component (48 hours pre, during, and post event).

Soil O_2_ at 30 cm decreased in both the freshwater and saltwater plots during the event, reaching anoxia toward the end of the event in both treatment plots (Figs 2 and 3). Interestingly, microbial CO_2_ fluxes in the root exclusion subplots increased noticeably in the saltwater plot early on during the TEMPEST flooding event, resulting in a positive change during the event (Fig 2), and decreased toward the end of the event in both plots (Fig S3).

The largest difference in biogeochemical responses between the freshwater and saltwater treatments was porewater DOC, which decreased in the saltwater plot and to a lesser extent in the freshwater plot (Fig 2). However, the trends in soil porewater DOC concentrations were similar in both treatment plots and the control plot immediately following the flooding event (Fig S3). The spectral slope ratio (Sr), a metric of porewater DOC quality, responded weakly (i.e. within control plot variability) but in opposite directions, decreasing in the freshwater plot and increasing in the saltwater plot (Fig 2). Microbial CH_4_ flux responded weakly relative to variance in the control plot and in different directions, with a decrease in the freshwater plot and an increase in the saltwater plot (Fig 2). The microbial fluxes from the saltwater plot root exclusion subplots momentarily shifted from being a methane sink (i.e., negative flux) to a source (i.e., positive flux to the atmosphere) during the flood treatment, whereas the root exclusion subplots in the freshwater and control plots remained CH_4_ sinks throughout the experiment (Fig S3). Finally, groundwater dissolved O_2_ responses were lagged, noticeably increasing after the flooding event, and remaining elevated in the freshwater plot during the 48 hours following the event (Fig S3).

### Vegetation responses

Vegetation responses to our short-term treatments were not uniform across the treatment plots. Root respiration in the saltwater plot and tree sap flow velocity in both treatment plots exhibited were the only measured vegetation variables that exhibited changes outside the control plot variance.

Root-influenced CH_4_ fluxes measured in the root exclusion subplots increased in the freshwater plot and decreased slightly in the saltwater plot (Fig 2), though these did not exceed the envelope of control plot variance. In the freshwater plot, root-influenced CH_4_ fluxes shifted from an atmospheric sink (i.e., negative flux) to an atmospheric source during the flood treatment whereas the saltwater plot shifted to a source only momentarily during and after the treatment (Fig 3). The change for the root CO_2_ fluxes was lower in both plots (Fig 2); however, this was only the saltwater plot was outside of the envelope of control plot variance.

Root-influenced CH_4_ fluxes peaked during the event in the freshwater plot, after the flooding event in the saltwater plot (Fig 3). Root respiration was lowest during the flooding event in both plots, but strong conclusions cannot be drawn because of high variation (Fig S4). Evaluating temporal patterns in vegetation responses, tree sap flow velocity followed consistent diel patterns throughout the flooding event (Fig S4).

### Natural rain event

During the rain event, volumetric water content at 15 cm depth increased in all plots (Fig 3). Soil O_2_ in both treatment plots remained lower than the pre-event values throughout the rain event and decreased in the control plot during the rain event (Fig 3). Groundwater dissolved O_2_ increased during the rain event in all plots (Fig S3). Tree sap flow velocity was lower in all plots during the rain event (Fig S4).

## Discussion

The punctuated hydrologic disturbance we applied to the ecosystem was the same order of magnitude as a tropical storm or hurricane in the region [21], and the increase in groundwater level observed in response to this event is analogous to increases in water table observed following hurricanes along the East Coast of the US [35]. Likewise, the increases we observed in groundwater salinity in the saltwater plot has been previously observed in coastal upland forests following a hurricane event [36]. We consider our attempt to simulate the freshwater and saltwater hydrologic responses to a hurricane-scale event successful. In observational studies of ecosystem responses to natural hurricane events, the contributions to tree mortality of aboveground impacts such as crown damage versus belowground impacts such as flooding or salt stress are difficult to disentangle [37]. Therefore, to remove these potentially confounding mechanisms, our experimental approach isolates the subset of mechanisms that operate primarily through belowground biogeochemical perturbations to advance process-level knowledge and numerical models capable of forecasting the longer-term impacts of flooding and salinity exposure in upland forest ecosystems.

The single novel flooding events simulated in the first application of TEMPEST significantly perturbed several hydrologic parameters but, as expected, the effects were transient. We hypothesized the hydrologic effects of our first freshwater and saltwater treatments would be the same despite the potential for salts to alter soil properties [38]. This was supported by the rapid decline in soil electrical conductivity in the 48 hours following the event across the saltwater plot. Although similar transient effects have been reported in other ecosystem-scale pulse saltwater addition experiments [38], the rain event that occurred less than 24 hours after treatments ended likely accelerated the decline in soil conductivity by flushing out the added ions from the treatment waters (Fig 4). In the absence of a rain event we expect that electrical conductivity (a proxy for salinity) would have remained elevated for a longer period of time [24,39]. Other studies have found no significant effect of freshwater additions on soil electrical conductivity [40,41], however, we attribute the freshwater increase in soil electrical conductivity to increase in electromigration due to increasing soil water content and not necessarily increase in salinity. Also, it is worth noting that the municipal freshwater source used for the experiment in relation to pre-event soil conductivity has a slightly elevated electrical conductivity (127 µS/cm) which contributed to increase in the measured electrical conductivity. This elevated electrical conductivity was far too low (0.06 PSU) to cause stress effects, however (< 0.5 PSU is generally considered to be freshwater [42]).

Even transient changes in soil saturation and salinity can perturb soil biogeochemical processes and plant physiology through shifts in soil O_2_ and changes in soil ionic strength [35,37]. The transient response of soil and groundwater salinity and O_2_ observed here may explain the lack of large or sustained shifts in biogeochemical and vegetation variables to the novel treatments. Despite over half of the variables examined exceeding variability in the control plot (7 of 13), the responses were generally either inconsistent in direction or small in magnitude. The lack of systematic effects may reflect high within-variable variation, relatively high variability in the control plot, or both (Figs 3 and S2-4). However, it more likely reflects the brief (nine hour) length of these pulse disturbances. This is consistent with the results of freshwater tidal marsh experiments where novel exposure to saline water added as a pulse caused relatively weak biogeochemical and vegetation responses compared to press disturbances [38,40,43]. Many soil flooding events that occur during and after natural hurricanes can last much longer than our event, on the order of days, leading to longer-lived changes in soil O_2_ and salinity, and stronger impacts on biogeochemical cycles [44]. The subtle differences in biogeochemical and vegetation responses between the freshwater and saltwater plots in the present study may be early indicators of stronger responses as the duration of exposure to flooding and salinity treatments is extended in future years of the TEMPEST experiment.

The relatively subtle and/or transient responses in focal hydrological, biogeochemical, and vegetation variables suggest this upland forest ecosystem is resistant to a single extreme exposure to fresh or brackish water. However, in anticipation of our plans to increase the frequency and duration of flood events, it is useful to consider whether the direction of the responses provide insights into mechanisms that will eventually drive more dramatic responses [20,22,23], such as our expectation of more persistent increases in soil conductivity discussed earlier. For example, the large decline in porewater DOC in the saltwater plot compared to the freshwater plot was likely driven by estuarine salts reducing carbon solubility in porewater solutions [45], an effect that is expected to impact microbial activity, soil organic matter decomposition, greenhouse gas (GHG) fluxes, and hydrologic export of DOC to the adjacent estuary if salts accumulate in the plot’s soils [45–47]. DOC chemical composition based on spectral slope responded very weakly but in different directions, increasing slightly in the saltwater plot, suggesting saltwater will preferentially remove higher molecular weight compounds from the DOC pool while freshwater flooding may have the opposite effect [48]. The relative decline in soil O_2_ was surprising for a short-term event and suggests that this upland forest, with typically aerobic soils, may be capable of developing quantitatively important anaerobic biogeochemical cycles quite rapidly as the duration of events lengthens from several hours to several days [49].

The long-term implications of changes in CO_2_ and CH_4_ fluxes are harder to discern from short-term responses alone because they are determined by interacting physical and biological factors. The brief increase we observed in root-free (heterotrophic) soil CO_2_ emissions has been observed in previous studies of soil respiration (heterotrophic + autotrophic) and may be due to physical displacement of CO_2_ in the soil matrix [50,51]. CH_4_ uptake by soils also responds to physical factors such as soil moisture content that control CH_4_ diffusion from the atmosphere [52–54]. Such physical factors are likely to dominate GHG responses over short time scales (hours), while biogeochemical and vegetation response become increasingly important as the frequency or duration of events increase [55]. Changes in soil CO_2_ emissions following hurricanes are rarely measured and the responses vary [40,56–59]. Further field observations coupled with lab experiments are needed to isolate the mechanisms that distinguish short-term from long-term responses to freshwater and saltwater flooding events.

We hypothesized that hydrology, biogeochemistry, and vegetation variables will respond at different rates to a novel flooding event, with vegetation responding most slowly. We chose to evaluate three vegetation-related variables that are most likely to respond within our 48 hour response window – tree sap flow velocity, root respiration, and root-influenced soil CH_4_ flux – understanding that the influence of roots on CH_4_ flux would operate indirectly through physical or microbial processes. Sap flow responded in opposite directions in the two treatments. Freshwater addition increased sap flow suggesting that transpiration was water-limited, a possibility that aligns with evidence of water limitation of forest NPP at the site [60]. The decrease in sap flow when the added water was saline suggests a stress response [20]. The decrease in root respiration in the freshwater and saltwater plots suggests a stress response that may have reduced root respiration rates [61,62]. However, the large variation in root respiration within each plot (Fig S4) argues for additional observations to establish cause and effect mechanisms.

Transient changes in soil moisture and oxygen levels such as those observed in response to our first brief TEMPEST treatment can influence long-term responses by changing baseline conditions, affecting the ability of the system to respond to future perturbations. The effect of the TEMPEST treatment on antecedent conditions is apparent in the response of the system to a rain event that occurred < 24 hours after the treatments ended. Soil O_2_ concentrations in the saltwater plot recovered quickly to about half of atmospheric saturation before rain began, while concentrations in the control plot started at full saturation. As a result of this difference in initial conditions, soil O_2_ concentrations were depleted much faster in the saltwater than control plots. The fact that soil O_2_ concentrations in the freshwater plot never recovered between the treatment and rain events reflects differences in the hydrology of the two plots that also affected antecedent conditions. The brief anaerobic conditions that occurred during the flooding event could set up antecedent changes in belowground redox, carbon, and nutrient cycling that may become more pronounced with repeated episodic events [63,64] in comparison to continuous flooding [65] inducing shifts in key processes regulating vegetative dynamics leading to plant stress and mortality [20].

Long-term, ecosystem-scale manipulations such as TEMPEST are vital for understanding and predicting ecosystem stress, change, and transformation through time [66–68]. Soil transplant and mesocosm studies find that antecedent conditions effectively regulate belowground biogeochemical responses to flooding and salinity [69–72]. In particular, sites experiencing infrequent or new exposure to salinity may be more sensitive than sites with more frequent exposures [69]. Previous studies in systems with episodic historical exposure to salinity are consistent with this idea, finding that chronic salinity exposure caused the most changes in ecosystem functions, such as GHG production, and the acute (press) treatments did not result in any sustained shifts [43]. This highlights the importance of understanding novel exposure, even when immediate responses are relatively muted and do not induce rapid state change. As coastal upland ecosystems undergo transitions to more flooding- and salinity-dominated systems, the trajectory of this transition likely also depends on starting system conditions. Differences in soil, vegetation, and hydrology in upland forests, compared to coastal freshwater wetland systems, may influence the trajectories of how upland forests respond to flooding and salinity. For example, shifts in soil carbon solubility and microbial respiration are linked to initial soil pH and composition [45], while soil salinity dynamics may be tied to initial soil moisture.

The ecosystem-level analysis we present serves as a baseline to understand acute belowground-driven shifts in ecosystem responses to flooding and salinity. The interactions among short-term salinity pulses, long-term hydrologic shifts, and vegetative responses to compounding disturbances can influence belowground carbon budgets in coastal marsh ecosystems [56], and may be early drivers of ecosystem state change in coastal upland forests. Outcomes from the novel, ecosystem-scale TEMPEST experiment provide information on the drivers, responses, and mechanisms of the impacts of hydrologic disturbance events on short- and long-term trajectories. Such information will be crucial to understand and predict changes in coastal ecosystem structure and function driven by changing environmental conditions expected throughout the 21st century [68].

## Acknowledgements

We thank the massive field sampling and infrastructure teams that made this first TEMPEST event possible – in particular, Rick Smith, Stella Woodard and Evan Phillips. We also thank James Stegen for his contributions to early conceptualization of the TEMPEST experimental design. This research was supported by COMPASS-FME, a multi-institutional project supported by the U.S. Department of Energy, Office of Science, Biological and Environmental Research as part of the Environmental System Science Program, and by the Smithsonian Environmental Research Center. The Pacific Northwest National Laboratory is operated for DOE by Battelle Memorial Institute under contract DE-AC05-76RL01830.

## TEMPEST 1.0 event consortium authors

Rick Smith, Anya Hopple, Pat Megonigal, Alice Steans, Evan Phillips, Stella Woodard, Stephanie Pennington, Ben Bond-Lamberty, Nate McDowell, Gary Peresta, Kendal Morris, Donnie Day, Wei Huang, Mia DiCianna, Lani DuFresne, Sam Wright, Allison Myers-Pigg, Opal Otenburg, Khadijah Homolka, Madison Bowe, Nicholas Ward, Roberta Peixoto, Jonathan Kwong, Peter Regier, Elaine Yu, Kennedy Doro, Moses Adebayo, Emmanuel Efemena, Solomon Ehosioke, Mitchell Smith, Yerang Yang, Drew Peresta, J. Alan Roebuck Jr., Leticia Sandoval

## Author contributions

Conceptualization: ANMP, AH, SP, PR, BBL, KD, NM, NW, VB, PM

Data Curation: ANMP, AH, SP, PR, MD, KD, JM

Formal Analysis: ANMP, SP, PR, MD, JM Funding Acquisition: BBL, NW, VB, PM

Investigation: TEMPEST 1.0

Event Consortium Methodology: TEMPEST 1.0 Event Consortium

Project Administration: ANMP, AH, SP, BBL, KD, NM, AS, NW, VB, PM

Resources: ANMP, BBL, KD, NM, NW, VB, PM

Software: ANMP, AH, SP, PR, JM

Supervision: ANMP, AH, SP, BBL, KD, NM, NW, PM

Validation: ANMP, AH, SP, PR Visualization: ANMP, SP, PR, AS

Writing – Original Draft: ANMP, SP, PR, PM

Writing – Review & Editing: ANMP, AH, SP, PR, BBL, MD, KD, NM, JM, AS, NW, VB, PM

## Supporting information captions

**Fig S1.** Plot installation locations.

**Fig S2.** Hydrological variable responses (soil conductivity, groundwater depth, groundwater conductivity) through time.

**Fig S3.** Biogeochemical variable responses (porewater DOC, microbial CH_4_ flux, microbial CO_2_ flux, porewater slope ratio, and groundwater DO) through time.

**Fig S4.** Vegetation variable responses (sap flow velocity, root carbon dioxide flux) through time.

